# Rapid and automated quantification of TDP-43 and FUS mislocalisation for screening of frontotemporal dementia and amyotrophic lateral sclerosis gene variants

**DOI:** 10.1101/2021.03.07.433817

**Authors:** Lisa J. Oyston, Stephanie Ubiparipovic, Lauren Fitzpatrick, Marianne Hallupp, Lauren M. Boccanfuso, John B. Kwok, Carol Dobson-Stone

## Abstract

**Background:** Identified genetic mutations cause 20% of frontotemporal dementia (FTD) and 5-10% of amyotrophic lateral sclerosis (ALS) cases: however, for the remainder of patients the origin of the disease is uncertain. The overlap in genetic, clinical and pathological presentation of FTD and ALS suggests these two diseases are related. Post-mortem, 97% of ALS and ∼50% of FTD patients show redistribution of the nuclear proteins TDP-43 or FUS to the cytoplasm within affected neurons. We exploited this predominant neuropathological feature to develop an automated method for the quantification of cytoplasmic TDP-43 and FUS in human cell lines.

**Results:** Utilising fluorescently-tagged cDNA constructs to identify cells of interest, the fluorescence intensity of TDP-43 or FUS was measured in the nucleus and cytoplasm of HEK293 and SH-SY5Y cells. Confocal microscope images were input into the freely available software CellProfiler, which was used to isolate and measure the two cellular compartments. Significant increases in the amount of cytoplasmic TDP-43 and FUS were detectable in cells expressing known ALS-causative *TARDBP* and *FUS* gene mutations. Pharmacological intervention with the apoptosis inducer staurosporine also induced measurable cytoplasmic mislocalisation of endogenous FUS. Additionally, this technique was able to detect the subtler effect of mutation in a secondary gene (*CYLD*) on endogenous TDP-43 localisation.

**Conclusions:** These findings validate this methodology as a novel *in vitro* technique for the quantification of TDP-43 or FUS mislocalisation that can be used to assess the pathogenicity of predicted FTD- or ALS-causative mutations.

## Background

Frontotemporal dementia (FTD) is one of the most common forms of presenile dementia and involves the progressive degeneration of the frontal and temporal lobes of the brain. Amyotrophic lateral sclerosis (ALS) is a progressive neurodegenerative disorder that affects the upper and lower motor neurons leading to muscle weakness and paralysis [1]. Increasing genetic evidence, neuropathological and clinical observations have identified a substantial overlap between these two disorders [2]. Several genes have been identified where mutations can cause either FTD or ALS (FTD-ALS genes) including *C9orf72, VCP, OPTN, SQSTM1, TBK1* [3] and, most recently, *CYLD* [4]. In addition, FTD and ALS share neuropathological similarities: ∼95% of ALS and ∼50% of FTD patients show cytoplasmic inclusions of the proteins TAR DNA-binding protein 43 (TDP-43) [5, 6] or fused in sarcoma (FUS) within the brain (∼5% of ALS and ∼10% of FTD patients) [6–8]. In healthy cells, TDP-43 and FUS are largely expressed within the nucleus. In FTD and ALS, affected neurons predominantly display a redistribution of TDP-43 or FUS to the cytoplasm, as well as insoluble TDP-43/FUS aggregates [6, 9, 10]. The importance of TDP-43 and FUS is further demonstrated by the fact that mutation of their encoding genes *TARDBP* and *FUS* is sufficient to cause ALS [7, 11, 12] or, rarely, FTD [13].

Identified genetic mutations cause 20% of familial FTD and 5-10% of familial ALS cases [14, 15]; however, for the remainder of patients the origin of the disease is uncertain. Despite TDP-43 and/or FUS mislocalisation being a prominent feature in the majority of FTD and ALS cases, including all those carrying FTD-ALS gene mutations [13], this fact has not been applied to the screening of FTD and ALS candidate gene variants on a large scale. In previous studies, TDP-43 and FUS mislocalisation has largely been assessed by subcellular fractionation, immunohistochemistry and manual analysis of confocal microscopy [7, 16–24]. Whilst studies utilising these techniques have been informative, these assays are labour intensive, low-throughput, expensive and often produce qualitative data. In addition, microscopy-based techniques could be subject to bias due to manual selection of the cells to be analysed.

Here we report the development of a rapid and automated technique for the *in vitro* study of TDP-43 and FUS cytoplasmic mislocalisation. TDP-43 or FUS staining in nuclear and cytoplasmic subcellular compartments was measured in thousands of cells from confocal microscope images, using the freely available analysis software CellProfiler [25]. We validated this technique using known genetic and pharmacological drivers of TDP-43 or FUS mislocalisation. We were able to detect increased cytoplasmic localisation of exogenously expressed *FUS* and *TARDBP* mutations relative to wild-type (WT) sequence. We also observed mislocalisation of endogenous FUS upon treatment with the apoptosis inducer staurosporine. Importantly, we could also detect more subtle changes in the cellular distribution of endogenous TDP-43, arising from mutation in a secondary gene (*CYLD* p.M719V). This validated methodology can now be utilised for rapidly assessing the pathogenicity of newly discovered FTD and ALS gene variants by quantifying their effect on TDP-43 and/or FUS localisation.

## Results

### Detection of FUS cytoplasmic mislocalisation with exogenous expression of FUS mutations

In order to validate our method for the unbiased quantification of FUS and TDP-43 mislocalisation we initially utilised known pathogenic *FUS* mutations that drastically alter FUS subcellular localisation [7, 26]. Expression of green fluorescent protein (GFP)-tagged FUS protein with missense mutation R521C (FUS_R521C_) or truncation mutant FUS_R495X_ in the neuronal-like neuroblastoma cell line, SH-SY5Y, led to a significant increase in the fluorescence intensity of cytoplasmic FUS (**Fig. 1e-k**) when compared to FUS_WT_ (**Fig. 1a-c, h**). This increase in cytoplasmic FUS was accompanied by a corresponding decrease in the fluorescence intensity of nuclear FUS (**Fig. 1d**). Together, these resulted in a 9.5- and 12.5-fold increase in the FUS cytoplasmic/nuclear ratio for cells expressing FUS_R521C_ (0.97±0.04; p<0.0001) and FUS_R495X_ (1.28±0.04; p<0.0001), respectively, relative to FUS_WT_ (0.10±0.00; **Fig. 1l**). Similar results were found when overexpressing the FUS_R521C_ (1.04±0.05; p<0.0001) and FUS_R495X_ (1.81±0.10; p<0.0001) mutants in human embryonic kidney (HEK293) cells (**Fig. S1**), although the decrease in nuclear fluorescence intensity of FUS was not as prominent (**Fig. S1d**).

**Fig. 1.**
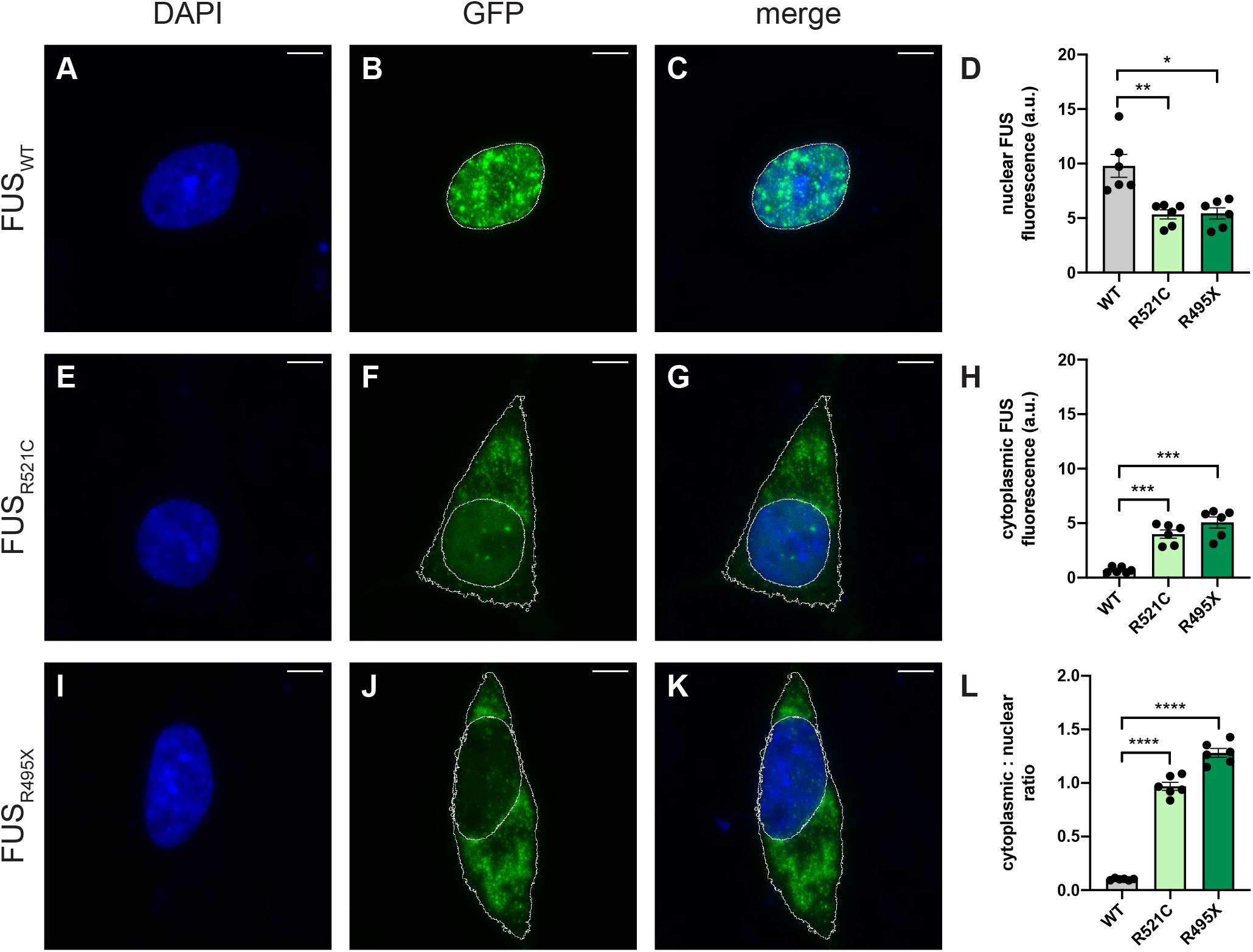
Detection of FUS cytoplasmic mislocalisation with exogenous expression of *FUS* mutations. Representative images of SH-SY5Y cells overexpressing GFP-tagged (**a-c**) FUS_WT_, (**e-g**) FUS_R521C_ or (**i-k**) FUS_R495X_. Nuclei were visualised with DAPI (blue). (**d**) Quantification of the fluorescence intensity of the nucleus shows a signifcant reduction in nuclear FUS expression in cells expressing FUS_R521C_ or FUS_R495X_, when compared to FUS_WT_. (**h**) Quantification of the fluorescence intensity of cytoplamic FUS shows significantly higher cytoplasmic FUS expression in FUS_R521C_ and FUS_R495X_, when compared to FUS_WT_. (**l**) Quantification of the cytoplasmic/nuclear ratio of exogenous FUS shows a marked increase in FUS_R521C_- and FUS_R495X_-expressing cells when compared to FUS_WT_. Scale bars = 5 µm. Data is represented as mean ± SEM. a.u. = arbitary units. *p<0.05; **p<0.01; ***p<0.001; ****p<0.0001.

### Detection of endogenous FUS cytoplasmic mislocalisation following staurosporine treatment

To test our ability to quantify the subcellular distribution of endogenous FUS we utilised the apoptosis inducer staurosporine, which has been previously reported to cause cytoplasmic mislocalisation of FUS [16]. Application of increasing doses of staurosporine showed an increase in the fluorescence intensity of cytoplasmic FUS in cells treated with the two highest dose of staurosporine (1 and 10 µM; **Fig. 2c-d, f**). A corresponding decrease in the nuclear intensity of FUS was also seen in SH-SY5Y cells (**Fig. 2e**). There was a significant increase of cytoplasmic/nuclear FUS ratio for all treatment groups (0.1 µM: 0.13±0.01; p=0.0405; 1 µM: 0.31±0.06; p=0.0152; 10 µM: 1.35±0.07; p=0.0002) relative to untreated cells (0.09±0.00; **Fig. 2g**). HEK293 cells treated with staurosporine also exhibited a dose-response effect on FUS subcellular localisation. In general, HEK293 cells were more tolerant of the staurosporine treatment (**Fig. S2a-d**) as shown by the lower cytoplasmic/nuclear ratios observed when compared to the SH-SY5Y cells (**Fig. 2g** and **S2g**). Treatment with 10 µM staurosporine caused the majority of FUS to be present in the cytoplasm rather than the nucleus in SH-SY5Y cells (1.35±0.07; p=0.0002; **Fig. 2d, g**) whilst the same treatment in the HEK293 cells caused only a small proportion of FUS protein to be mislocalised (0.28±0.01; p<0.0001; **Fig. S2d, g**). Despite these differences in effect size, we could detect a significant effect of staurosporine treatment on mislocalisation of endogenous FUS in both cell lines.

**Fig. 2.**
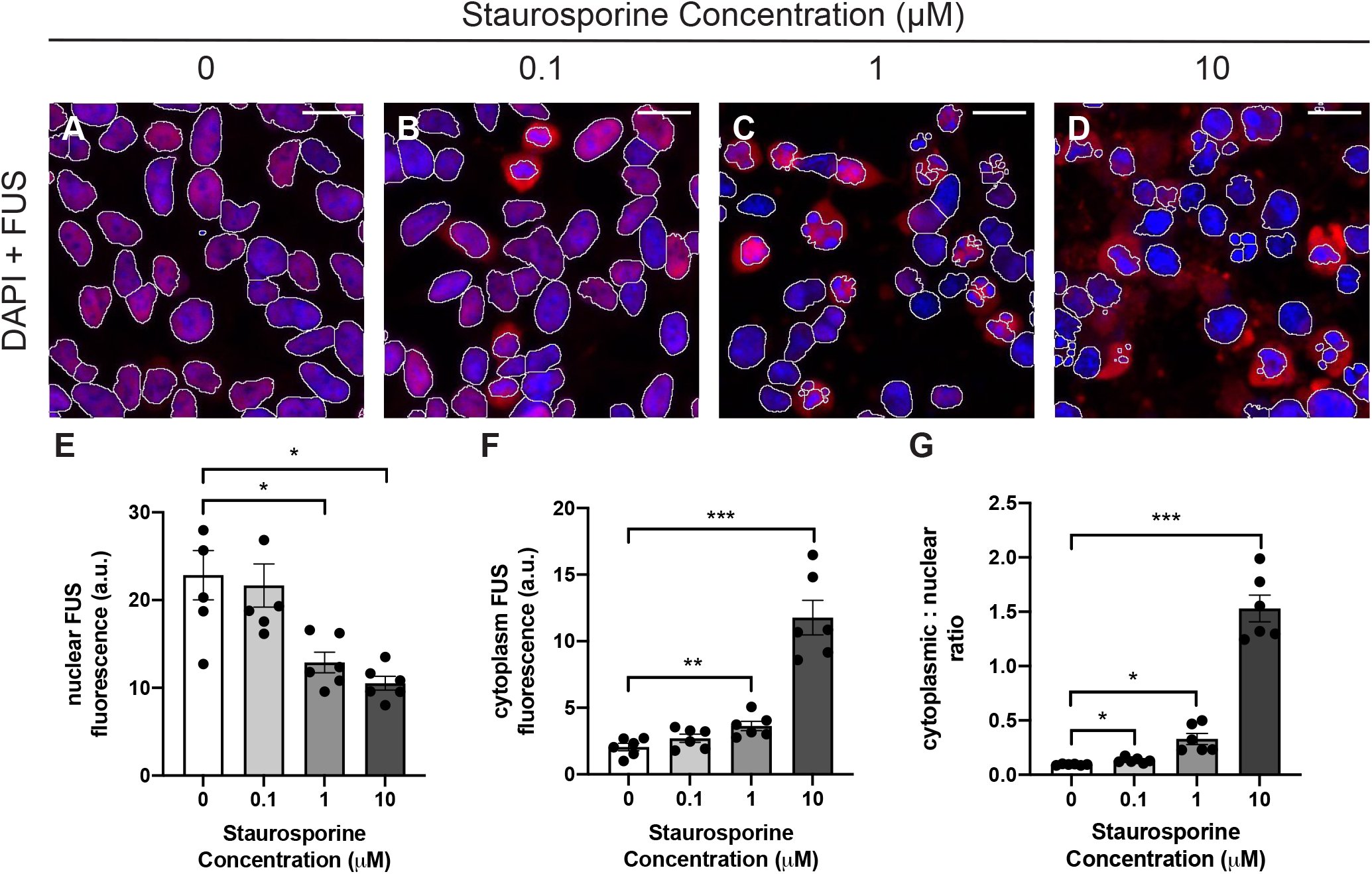
Detection of endogenous FUS cytoplasmic mislocalisation following staurosporine treatment. (**a-d**) Representative fluorescence images of SH-SY5Y cells treated with increasing concentrations of staurosporine. FUS was dectected by immunofluorescent staining (red) and nuclei were visualised with DAPI (blue). (**e**) Quantification of nuclear and (**f**) cytoplasmic FUS fluorescence intensity show significant changes in 1 µM- and 10 µM-treated cells. (**g**) Quantification of the FUS cytoplasmic/nuclear ratio shows an increase in all treatment groups when compared to those treated with DMSO alone. Scale bars = 20 µm. Data is represented as mean ± SEM. a.u. = arbitary units. *p<0.05; **p<0.01; ***p<0.001.

### Detection of TDP-43 cytoplasmic mislocalisation with exogenous expression of TARDBP mutations

We took the same initial approach used for FUS to validate this method for the quantification of TDP-43 cytoplasmic mislocalisation: i.e. exogenous expression of known pathogenic mutations in *TARDBP*. We examined p.A315T, one of the earliest detected and thus most extensively examined mutations [27–29], and two of the mutations most commonly observed in ALS patients: p.M337V and p.A382T [11, 28–30].

In SH-SY5Y cells, a significant increase in the cytoplasmic/nuclear ratio of GFP-tagged TDP-43_A315T_ (0.87±0.01; p=0.0016; **Fig. 3b, f, k**) and TDP-43_A382T_ (0.86±0.02; p=0.0032; **Fig. 3d, h, k**) was observed when compared to TDP-43_WT_ (0.75±0.01; **Fig. 3a, e, k**). In contrast, there was a significant decrease in the cytoplasmic/nuclear ratio of GFP-tagged TDP-43_M337V_ (0.63±0.01; p=0.0028; **Fig. 3c, g, k**) relative to TDP-43_WT._ Similar to the experiments with FUS, expression of TDP-43 mutations had a lesser effect in HEK293 cells (**Fig. S3**). Once again, there was a significant increase in the cytoplasmic/nuclear ratio of TDP-43_A315T_ (0.57±0.02; p=0.0021; **Fig. S3b, f, k**) and TDP-43_A382T_ cells (0.66±0.03; p=0.0026; **Fig. S3d, h, k**) when compared to TDP-43_WT_ (0.50±0.02; **Fig. S3a, e, k**). However, no significant effect was observed in TDP-43_M337V_ cells (0.52±0.01; **Fig. S3c, g, k**).

**Fig. 3.**
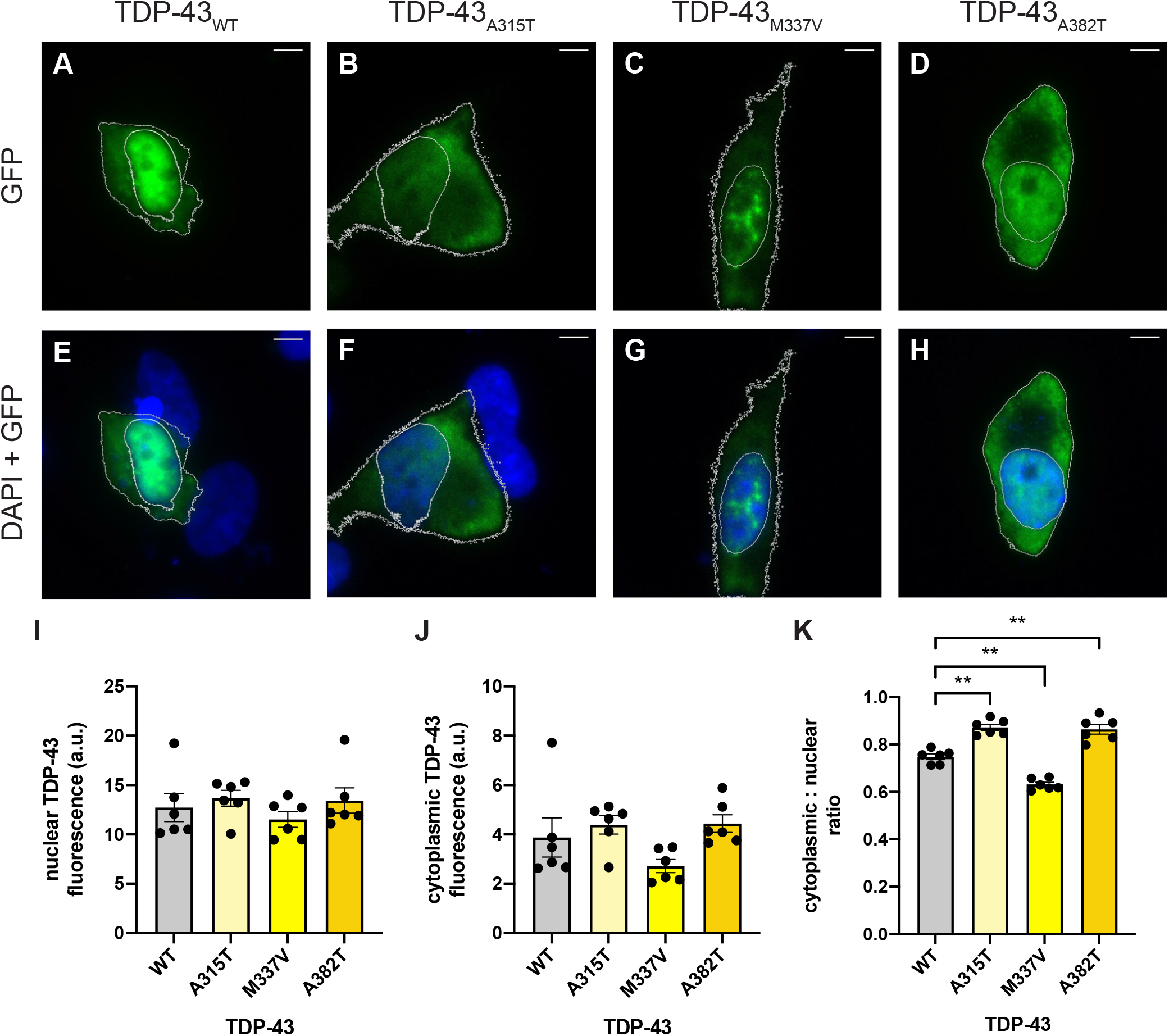
Detection of TDP-43 cytoplasmic mislocalisation with exogenous expression of *TARDBP* mutations. Representative images of SH-SY5Y cells overexpressing GFP-tagged (**a, e**) TDP-43_WT_, (**b, f**) TDP-43_A315T_, (**c, g**) TDP-43_M337V_ or (**d, h**) TDP-43_A382T_. Nuclei were visualised with DAPI (blue). (**i**) Quantification of the fluorescence intensity of the nucleus and (**j**) the cytoplasm shows no significant difference in TDP-43 expression. (**k**) Quantification of the cytoplasmic/nuclear ratio of exogenous TDP-43 shows a significant increase in TDP-43_A315T_ and TDP-43_A382T_, and a decrease in TDP-43_M337V_, when compared to cells expressing TDP-43_WT_. Scale bars = 5 µm. Data is represented as mean ± SEM. a.u. = arbitrary units. **p<0.01.

### Detection of endogenous TDP-43 cytoplasmic mislocalisation in cells expressing a causative FTD/ALS mutation

To demonstrate the potential of this methodology for evaluating the pathogenicity of variants in other FTD/ALS genes, we expressed a known familial FTD/ALS-causative mutation in the CYLD gene, p.M719V [4], and observed the changes in endogenous TDP-43 localisation. We previously determined that overexpression of CYLD_M719V_ increases the proportion of cytoplasmic TDP-43-positive mouse primary cortical neurons by ∼20% when compared to CYLD_WT_ [4]. When compared to GFP-only vector control (0.079±0.001; **Fig. 4a-d**), expression of CYLD_WT_-GFP (**Fig. 4e-h**) in SH-SY5Y cells led to a significant 1.2-fold increase in the TDP-43 cytoplasmic/nuclear ratio (0.095±0.001; p=0.0002; **Fig. 4s**). In turn, expression of CYLD_M719V_-GFP (**Fig. 4m-p**) caused a further 1.3-fold increase in cytoplasmic/nuclear TDP-43 relative to CYLD_WT_-GFP (0.121±0.002; p<0.0001). The CYLD_M719V_ mutation had the same effect in HEK293 cells, causing a 1.3-fold increase in the cytoplasmic/nuclear ratio relative to CYLD_WT_ (0.145±0.007; p=0.0115; **Fig. S4e-h, m-p, s**). We also transfected cells with the CYLD_D681G_ mutation (**Fig. 4i-l, Fig. S4i-l**) which is catalytically inactive and known to cause CYLD cutaneous syndrome, a skin tumour disorder [31, 32]. Our data corroborated previous results [4] that the CYLD_D681G_ mutation had no effect on TDP-43 localisation when compared to the GFP-only vector control in either cell type (**Fig. 4s, Fig S4s**).

**Fig. 4.**
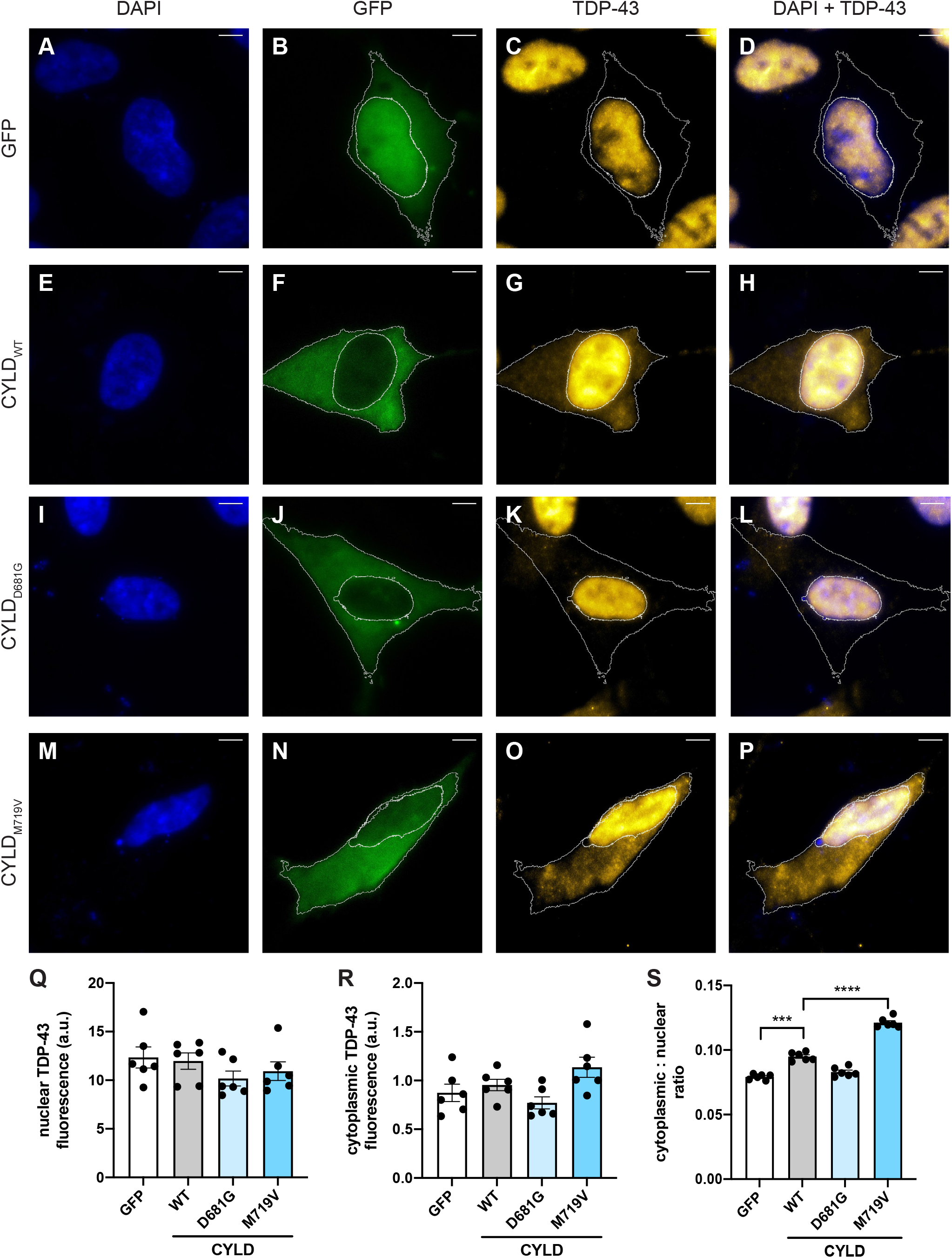
Detection of endogenous TDP-43 cytoplasmic mislocalisation in cells expressing *CYLD* mutations. Representative images of SH-SY5Y cells overexpressing (**a-d**) GFP or GFP-tagged (**e-h**) CYLD_WT_, (**i-l**) CYLD_D681G_ or (**m-p**) CYLD_M719V._ TDP-43 was detected by immunofluorescent staining (red) and nuclei were visualised with DAPI (blue). (**q**) Quantification of the fluorescence intensity of the nucleus shows no difference in TDP-43 expression. (**r**) Quantification of cytoplasmic TDP-43 fluorescence intensity shows increased TDP-43 expression in CYLD_WT,_ when compared to GFP and CYLD_M719V,_ when compared to CYLD_WT_. (**s**) Quantification of the cytoplasmic/nuclear ratio of exogenous TDP-43 shows a marked increase in CYLD_WT_, when compared to cells expressing GFP. The cytoplasmic/nuclear ratio of TDP-43 in CYLD_M719V_-expressing cells was significantly higher than CYLD_WT_. Scale bars = 5 µm. Data is represented as mean ± SEM. a.u. = arbitrary units. ***p<0.001; ****p<0.0001.

## Discussion

In this study we have validated a novel methodology for the automated quantification of cytoplasmic TDP-43 and FUS in human cell lines under three different experimental paradigms: determining localisation of exogenously expressed mutant TDP-43 or FUS; detecting the effect of a chemical modulator on endogenous FUS localisation; and detecting the effect of a secondary gene on endogenous TDP-43 localisation. This rapid, cheap, quantitative assay can now be utilised for rapidly assessing drug treatments and the pathogenicity of newly discovered FTD and ALS gene variants by quantifying their effect on TDP-43 and/or FUS localisation.

Previous studies examining TDP-43 and FUS have quantified cytoplasmic mislocalisation in different ways. Many studies have reported the proportion of the cell population that display cytoplasmic TDP-43 or FUS expression [12, 18, 19, 21, 24]. This may present a problem with reproducibility, since fluorescence detection thresholds for considering a cell as ‘positive’ for cytoplasmic protein likely differ between labs and will vary according to the microscopy equipment used. We selected the ratio of cytoplasmic to nuclear expression as our primary measure, having observed reduced variability between experimental replicates in comparison to individual nuclear and cytoplasmic measurements. Some studies have reported similar measurements using techniques such as high-content screening confocal microscopy to study TDP-43 or FUS mislocalisation under different conditions [33, 34]. Whilst high-content screening allows for a more high-throughput study design than our methodology, the extensive infrastructure required is expensive and can require substantial optimisation.

Another general observation in our experiments was that SH-SY5Y cells often displayed a greater degree of cytoplasmic mislocalisation than that in HEK293 cells under the same conditions. This may be due to SH-SY5Y cells being a neuronal cell line and thus more disease relevant, or due to the longer transfection time required to achieve optimal transfection efficiency in SH-SY5Y cells (48 hours, versus 24 hours in HEK293), causing additional stress on the cells. It should also be noted that transfection efficiency also dictated the number of images required to reach the desired cell number per replicate (300 cells). SH-SY5Y cells, which had a lower transfection efficiency require more images to achieve the same number of cells for quantification.

The cytoplasmic mislocalisation and/or aggregation of FUS in neurons and glia of post-mortem tissue from patients with FTD and ALS caused by FUS mutations has been widely established [7, 12, 35, 36]. This pathology has been recapitulated by expression of FUS mutations in mammalian cells [7, 12, 26]. In this study, exogenous expression of the severe truncation mutation FUS_R495X_ caused the majority of FUS to be mislocalised to the cytoplasm as previously demonstrated by Bosco et al. [26] using live-cell confocal microscopy and 3D images (11-28 cells). Our quick and simple methodology was able to duplicate these results using 2D images and a larger number of cells (1800 cells total). While we recognise the merits of quantifying the entire cell volume, measuring such a small number of selected cells is not representative of the population and may not be able to detect more subtle or variable effects in protein localisation.

Expression of the FUS_R521C_ missense mutation and treatment with staurosporine (1-10 µM) also reproduced previous qualitative microscopy and subcellular fractionation results [7, 16, 37] showing a significant increase in the FUS cytoplasmic/nuclear ratio. Confirmation of these results demonstrate the effectiveness of this new methodology in providing quantitative data where previously qualitative or labour-intensive experiments were required. Our ability to quantify FUS cytoplasmic mislocalisation following staurosporine treatment also demonstrates the possibility of this technique for use in testing the ability of novel FTD and ALS drug treatments to modulate FUS or TDP-43 mislocalisation.

The majority (∼95%) of ALS and ∼50% of FTD cases are characterised by the abnormal accumulation of TDP-43 in the cytoplasm of neurons and glia, even though in most cases mutations in *TARDBP* are absent [6]. Mutations in TDP-43 have been associated with both FTD and ALS and lead to TDP-43 cytoplasmic mislocalisation [11, 13]. Unlike FUS, the expression of TDP-43 mutations in primary cells and cell lines does not reproduce the dramatic cytoplasmic mislocalisation seen in post-mortem tissue [38–41]. In addition, overexpression of exogenous TDP-43 is sufficient to induce mislocalisation even for WT sequence [19, 23] Any differences between WT and mutant TDP-43 for this characteristic are therefore harder to discern in exogenous expression experiments and may be responsible for differing results being reported for the same mutations. Our technique was able to detect a significant increase in the TDP-43 cytoplasmic/nuclear ratio in TDP-43_A315T_ and TDP-43_A382T_ cells when compared to TDP-43_WT_. This confirms previous results for TDP-43_A315T_ [18, 19] and TDP-43_A382T_ [42, 43]. The TDP-43_M337V_ variant, which displayed a significant decrease in cytoplasmic TDP-43 in SH-SY5Y cells, has previously been shown to increase cytoplasmic TDP-43 in some [19, 21, 24], but not all studies [22]. We note that TDP-43_M337V_ has been shown to aggregate into nuclear puncta [21], which we observed in our experiments. As these puncta fluoresce at a higher intensity than diffuse TDP-43 expression, this may lead to an overall decrease in cytoplasmic/nuclear fluorescence ratio relative to TDP-43_WT_. Thus, in some cases quantification of nuclear and cytoplasmic puncta will be a useful adjunct to assess the pathogenicity of a given mutation. We also note that mislocalisation of TDP-43 is not the only mechanism by which *TARDBP* mutations lead to disease: for example, changes in protein stability or interaction partners have also been described [28]. However, we envisage that the mislocalisation assay as implemented here would be a useful tool in a battery of assays to screen novel *TARDBP* variants.

Lastly, we were able to recapitulate the effect of WT CYLD, the FTD-ALS-causative mutation p.M719V and the catalytically inactive mutation p.D681G on subcellular localisation of endogenous TDP-43, which we previously observed in mouse cortical neurons [4]. In comparison to the other experiments used to validate this quantification method, the degree of change in the endogenous TDP-43 cytoplasmic/nuclear ratio between the empty vector, the CYLD_WT_ and the CYLD mutations was very small: e.g., absolute increase in ratio of ∼0.03-0.04 between CYLD_WT_ & CYLD_M719V_ (**Fig. 4s**. and **S4s**). Our ability to detect significant differences for such a subtle change demonstrates the sensitivity of this technique for detecting changes in TDP-43 and FUS localisation *in vitro*. The utilisation of our quantification method in this format demonstrates what we believe to be its primary use, validating the pathogenicity of newly identified FTD- or ALS-associated variants in genes other than *FUS* and *TARDBP*.

## Conclusions

In summary, this study demonstrates a simple automated methodology for quantifying TDP-43 and FUS cytoplasmic mislocalisation *in vitro*. This technique can easily be added to studies utilising qualitative data to strengthen clearly noticeable phenotypes in an unbiased manner, as well as to highlight subtle changes that may have not been previously identified. Utilisation of this technique to aid in assessing pathogenicity of gene variants observed in patients with FTD or ALS will help to prioritise these variants for more intensive research efforts into how they cause disease.

## Methods

### DNA constructs

*CYLD, TARDBP* and *FUS* mutations were introduced into the pCMV6-CYLD, pCMV6-TARDBP and pCMV6-FUS constructs (Origene) by site-directed mutagenesis using the QuikChange Lightning Site-Directed Mutagenesis Kit (Agilent). Mutated *CYLD* and *TARDBP* cDNA sequences were then subcloned, using AsiSI and MluI restriction sites, into the pCMV6-AC-GFP vector (Origene) for expression of C-terminal GFP-tagged CYLD and TDP-43 proteins, respectively. Mutated *FUS* cDNA sequences were subcloned similarly into the pCMV6-AN-GFP vector (Origene) for expression of N-terminal GFP-tagged FUS protein, to avoid interference of the GFP tag with the C-terminal nuclear localisation signal of the FUS protein [44]. All clones were verified by restriction digestion and sequence analysis.

### Cell Culture

HEK293 and SH-SY5Y cell lines were used in this study. HEK293 cells were maintained in Eagle’s Minimum Essential Medium (EMEM; Gibco) and SH-SY5Y cells in Dulbecco’s Modified Eagle’s Medium/Nutrient Mixture F-12 (DMEM/F12; 1:1 mixture; Gibco) each containing 10% heat-inactivated fetal calf serum (Sigma-Aldrich). For immunocytochemistry, 8-well chamber slides (Ibidi) were coated with (concentration) poly-L-lysine (Sigma-Aldrich) for 1 h and then washed twice with Dulbecco’s phosphate-buffered saline (DPBS; Gibco). SH-SY5Y cells were seeded at 5– 7.5 × 10^4^ cells/well and HEK293 cells were seeded at 8 × 10^4^ cells/well. After 24 h, cells were transfected with GFP constructs (250 ng/well) for 24 (HEK293) or 48 (SH-SY5Y) hours using Lipofectamine 3000 (0.75 µL/well; Invitrogen) as per the manufacturer’s protocol. For the staurosporine experiment, cells were incubated for 48 h after seeding, then treated for 5 h with 0.1, 1 or 10 µM staurosporine (Sigma-Aldrich), to induce apoptosis. 100X staurosporine solutions were prepared from a 1 mM staurosporine stock solution in dimethyl sulfoxide (DMSO), thus DMSO was adjusted to a final concentration of 1% for control and lower staurosporine concentrations.

### Immunocytochemistry

Transfected and/or drug-treated cells were fixed in 4% paraformaldehyde (PFA) in PBS for 20 minutes in the dark. For visualisation of exogenous TDP-43 and FUS, cells were then washed twice with DPBS and mounted using DAKO fluorescent mounting medium containing 4’,6-diamidino-2-phenylindole (DAPI; Agilent). For visualisation of endogenous TDP-43 and FUS, non-specific background staining was blocked after cell fixation using 1% bovine serum albumin (BSA) in PBS for 30 minutes. Cells were then incubated overnight at 4°C with rabbit anti-TDP-43 (1:500; Proteintech #10782-2-AP) or rabbit anti-FUS (1:500, Proteintech #11570-1-AP) antibodies. The next day, cells underwent 2 x 5-minute washes with DPBS, incubation with rabbit Alexa Fluor-647 secondary antibody (1:500; #A-21244) for 1 hour, followed by 2 x 5-minute washes and mounting as above. Cells were imaged for fluorescence intensity quantification with a 40x objective on a Nikon A1R confocal microscope. Images of representative cells were obtained with a 100x objective for figure preparation. Laser intensity, gain and offset settings were kept constant and at least five random fields of view were imaged for each experiment.

### Fluorescence Intensity Quantification

Quantification of changes in the cellular localisation of TDP-43 or FUS between the nucleus and the cytoplasm was done by utilising CellProfiler 3.1.9 software (Broad Institute, Cambridge, MA) [25]. To prepare images for analysis, ND2 microscope files were converted to TIFFs, blinded and split into individual wavelength channel images using ImageJ (National Institutes of Health, Bethesda MD). In CellProfiler, the DAPI channel was analysed first using the *Identify Primary Objects* module and a global, three-class Otsu thresholding method. This allowed for identification and separation of each nuclei within an image. Strict filtering for cell diameter (30-60 pixels) eliminated any irregular or overlapped nuclei that were present. TDP- or FUS-positive cells were also identified using the *Identify Primary Objects* module. Cell diameter filtering for this step had a much larger range and was dependent on experimental conditions and cell type (20-150 pixels). The *Relate* and *Filter* modules were used to link filtered nuclei to TDP- or FUS-positive cells. Centroid distance filtering, which limits the distance between the centre of the nucleus and the centre of the cytoplasm (maximum 40 pixels) was also applied to remove any debris or abnormal cells that may be fluorescing in the TDP-43/FUS channel. For the experiments quantifying endogenous TDP-43 in cells expressing GFP-tagged CYLD constructs, additional steps in the analysis were required. GFP-positive CYLD cells were identified in the same way as TDP- or FUS-positive cells using the *Identify Primary Objects* module, described above. However, prior to this, TDP-positive cells were identified by association with their corresponding nuclei, using the *Identify Secondary Objects* module. The watershed-image method was used with global, three-class Otsu thresholding. Size filtering was also applied (maximum 3000 pixels) to remove any clumped cells. The *Relate* and *Filter* modules were used to select only TDP-positive cells that were also positive for GFP. TDP- and GFP-positive cells were then linked to their corresponding nuclei using the *Relate* and *Filter* modules described above. For all experiments, in each TDP-43/FUS-positive cell, the nucleus and cytoplasm were separated into individual objects using the *Identify Tertiary Objects* module. Integrated (total) fluorescence intensity for both cellular compartments were then measured from the TDP-43/FUS channel image using the *Measure Object Intensity* module. Using these measurements, the cytoplasmic to nuclear ratio of TDP-43 or FUS was calculated for 300 cells per group, for each of the six replicates of the experiment. All filtering and thresholding steps in each experimental pipeline were kept consistent throughout all experimental replicates. Filtering values were specific to each experiment and require adjustment for different cell types and microscope parameters.

### Statistical Analysis

Data are presented as mean ± standard error of the mean (SEM) from at least six independent experiments. Mean fluorescent intensity of TDP-43 or FUS across six biological replicates was in most cases compared by repeated measures one-way ANOVA and Dunnett’s multiple comparisons test. The effect of CYLD on endogenous TDP-43 was analysed using repeated measures one-way ANOVA and Sidak’s multiple comparisons test. All statistical analyses were performed using GraphPad Prism 9. Significance for all tests was set at p < 0.05.

## Abbreviations

ALS: amyotrophic lateral sclerosis
BSA: bovine serum albumin
DAPI: 4’,6-diamidino-2-phenylindole
DMEM: Dulbecco’s modified Eagle’s medium
DMSO: dimethyl sulfoxide
DPBS: Dulbecco’s phosphate-buffered saline
EMEM: Eagle’s minimum essential medium
FTD: frontotemporal dementia
FUS: fused in sarcoma
F12: nutrient mixture F-12
GFP: green fluorescent protein
HEK293: human embryonic kidney
PFA: paraformaldehyde
SEM: standard error of the mean
TDP-43: TAR DNA-binding protein 43
WT: wild-type

## Declarations

### Ethics approval and consent to participate

Not applicable.

### Consent for publication

Not applicable.

### Availability of data and materials

The datasets and materials generated during the current study are available from the corresponding author on reasonable request.

### Authors’ contributions

LJO, JBK and CDS conceived the study. SU, LF, MH and LMB generated and sequence-verified mutant cDNA constructs. LJO carried out TDP-43 and FUS localisation experiments. LJO and CDS participated in data analysis. LJO and CDS drafted the manuscript. All authors read and approved the final manuscript.

## Acknowledgements

The authors acknowledge the facilities and technical assistance of Microscopy Australia at the Australian Centre for Microscopy & Microanalysis at the University of Sydney.

