## Supplementary Figures for "Rapid and automated quantification of TDP-43 and FUS mislocalisation for screening of frontotemporal dementia and amyotrophic lateral sclerosis gene variants"

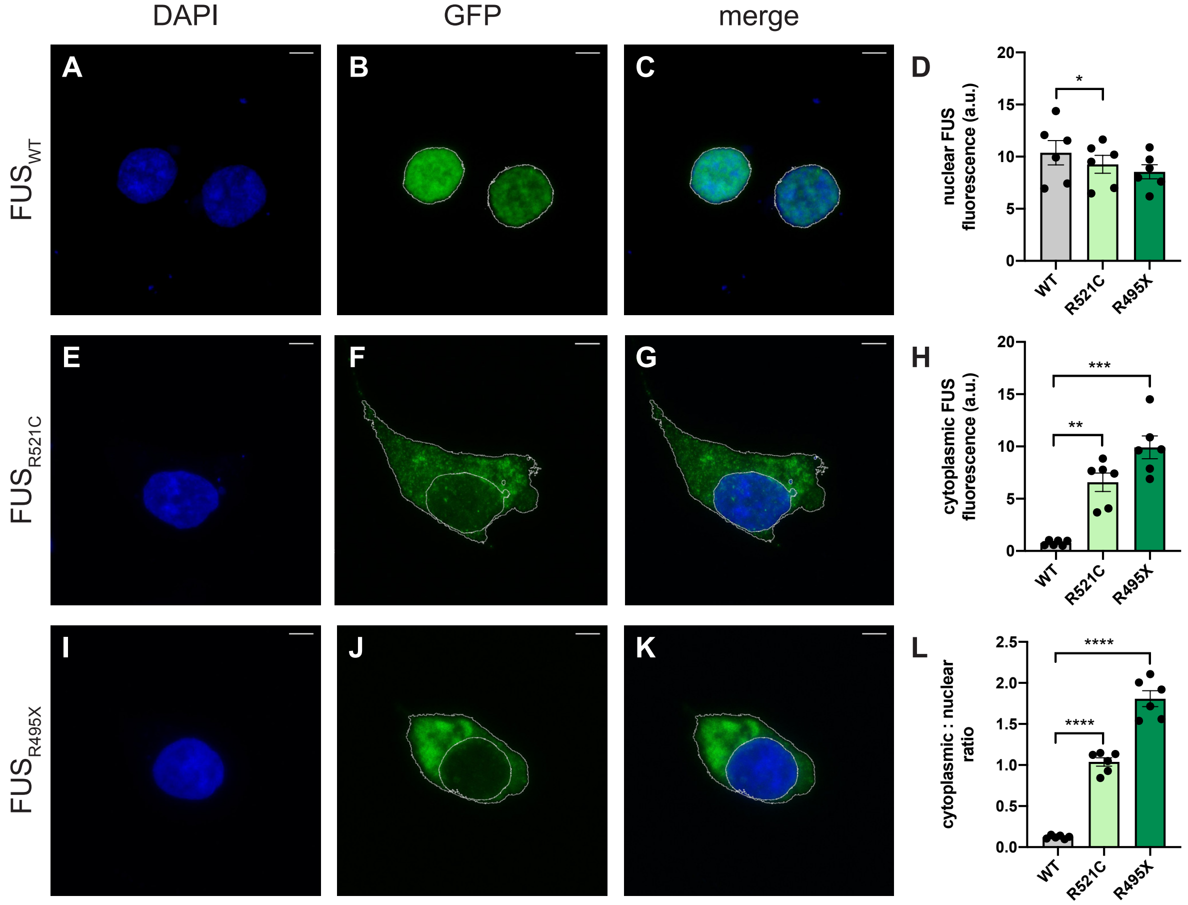


**Fig. S1 Detection of FUS cytoplasmic mislocalisation with exogenous expression of *FUS* mutations.** Representative images of HEK293 cells overexpressing GFP-tagged (**a-c**) $FUS_{WT}$, (**e-g**) $FUS_{R521C}$ or (**i-k**) $FUS_{R495X}$. Nuclei were visualised with DAPI (blue). (**d**) Quantification of the fluorescence intensity of the nucleus shows no difference in nuclear FUS expression. (**h**) Quantification of the fluorescence intensity of cytoplamic FUS shows significantly higher cytoplasmic FUS expression in $FUS_{R521C}$ and $FUS_{R495X}$, when compared to $FUS_{WT}$. (**l**) Quantification of the cytoplasmic/nuclear ratio of exogenous FUS shows a marked increase in $FUS_{R521C}$- and $FUS_{R495X}$-expressing cells when compared to $FUS_{WT}$. Scale bars = 5 µm. Data is represented as mean ± SEM. a.u. = arbitrary units. *p<0.05; **p<0.01; ***p<0.001; ****p<0.0001.

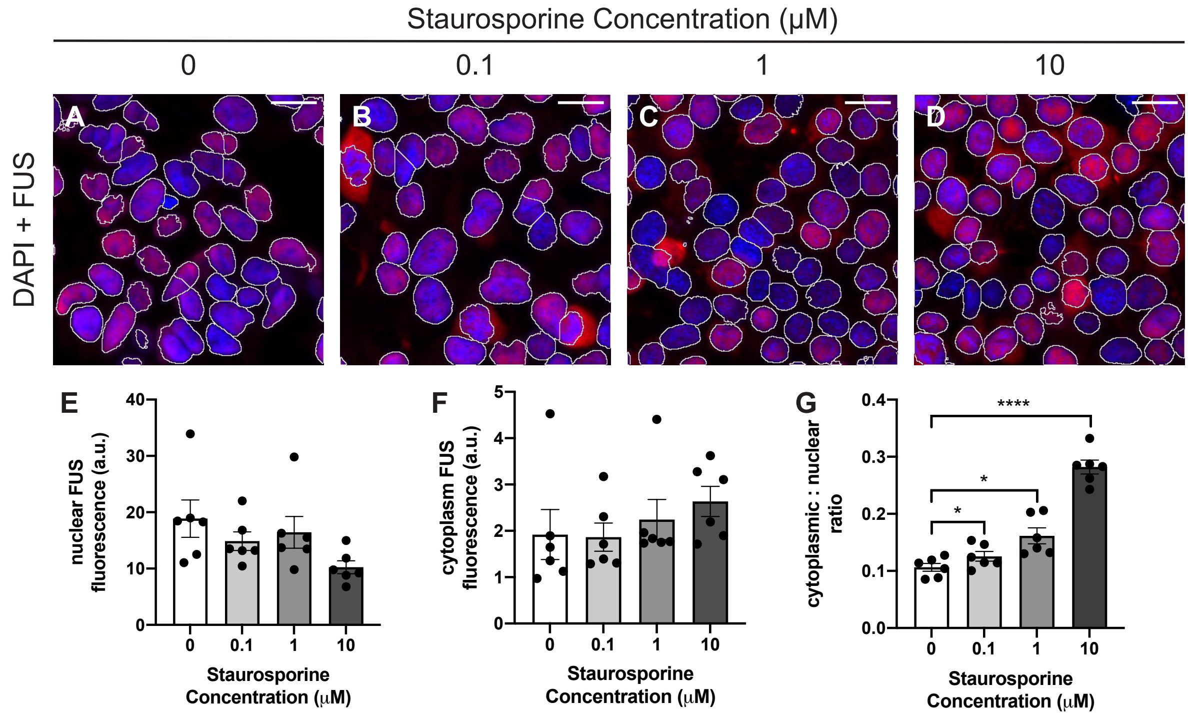


**Fig. S2 Detection of endogenous FUS cytoplasmic mislocalisation following staurosporine treatment.** (**a-d**) Representative fluorescence images of HEK293 cells treated with increasing concentrations of staurosporine. FUS was dectected by immunofluorescent staining (red) and nuclei were visualised with DAPI (blue). (**e**) Quantification of nuclear FUS fluorescence intensity and (**f**) cytoplasmic FUS fluorescence intensity showed no difference between groups. (**g**) Quantification of the FUS cytoplasmic/nuclear ratio showed an increase in all treatment groups when compared to those treated with DMSO alone. Scale bars = 20 μm. Data is represented as mean ± SEM. a.u. = arbitary units. *p<0.05; ****p<0.0001.

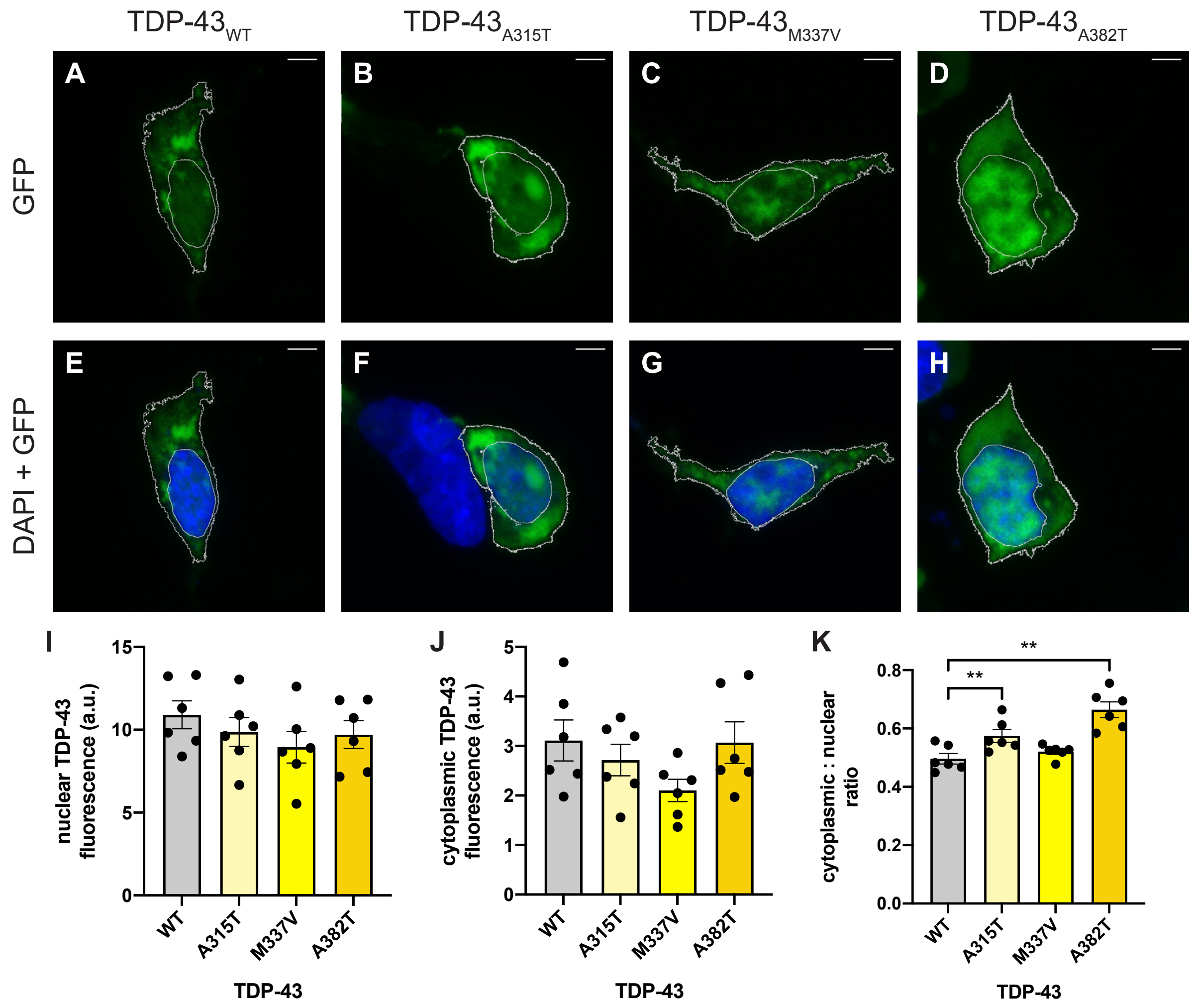


**Fig. S3 Detection of TDP-43 cytoplasmic mislocalisation with exogenous expression of *TARDBP* mutations.** Representative images of HEK293 cells overexpressing GFP-tagged (**a, e**) TDP-43$_{WT}$, (**b, f**) TDP-43$_{A315T}$, (**c, g**) TDP-43$_{M337V}$ or (**d, h**) TDP-43$_{A382T}$. Nuclei were visualised with DAPI (blue). (**i**) Quantification of the fluorescence intensity of the nucleus and (**j**) the cytoplasm shows no difference in TDP-43 expression. (**s**) Quantification of the cytoplasmic/nuclear ratio of exogenous TDP-43 shows an increase in TDP-43$_{A315T}$ and TDP-43$_{A382T}$-expressing cells, when compared to TDP-43$_{WT}$. Scale bars = 5 µm. Data is represented as mean ± SEM. a.u. = arbitrary units. **p<0.01.

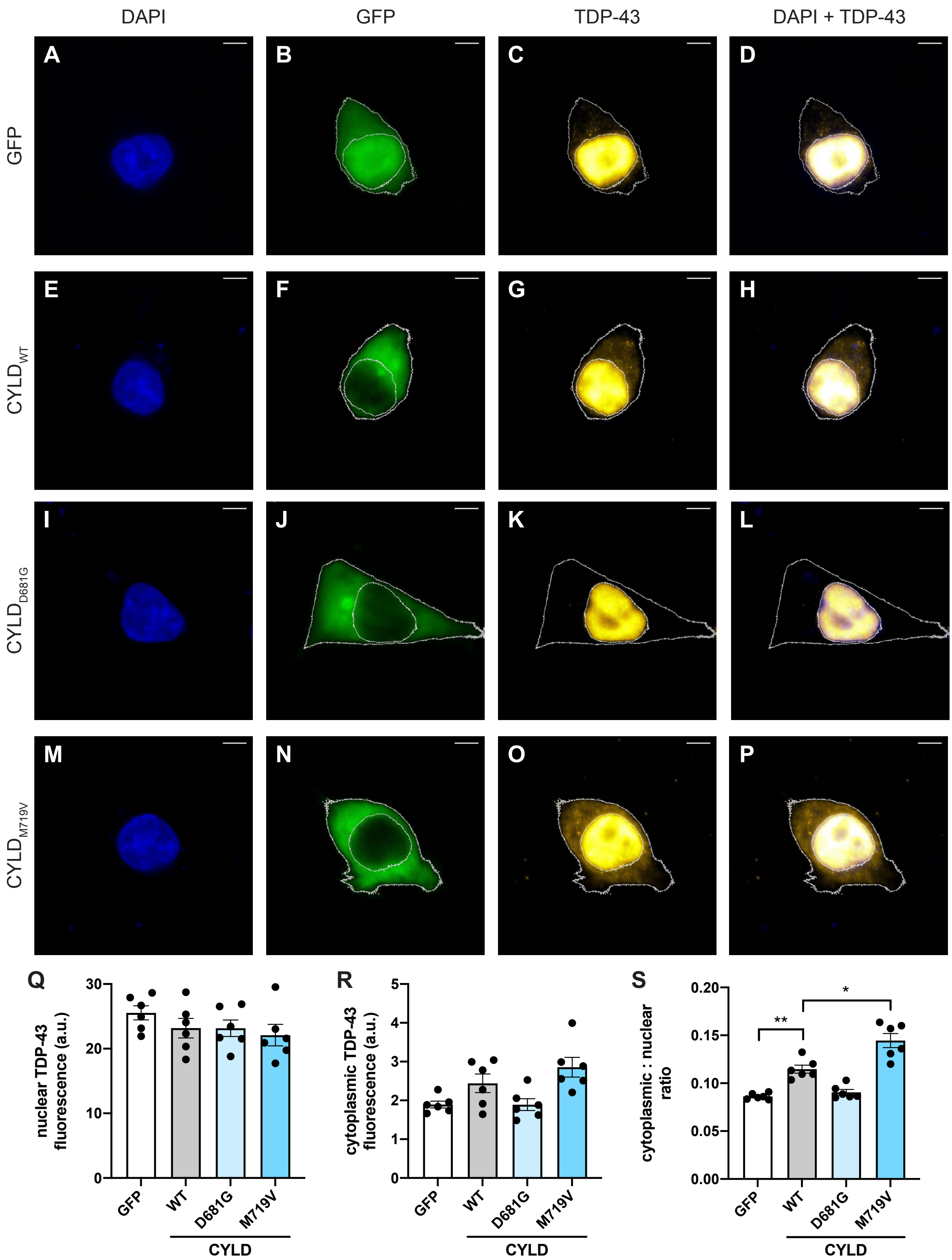


**Fig. S4 Detection of endogenous TDP-43 cytoplasmic mislocalisation in cells expressing *CYLD* mutations.** Representative images of HEK293 cells overexpressing (**a-d**) GFP or GFP-tagged (**e-h**) CYLD$_{WT}$, (**i-l**) CYLD$_{D681G}$ or (**m-p**) CYLD$_{M719V}$. TDP-43 was detected by immunofluorescent staining (red) and nuclei were visualised with DAPI (blue). (**q**) Quantification of the fluorescence intensity of the nucleus shows no difference in TDP-43 expression. (**r**) Quantification of cytoplasmic TDP-43 fluorescence intensity shows increased TDP-43 expression in $CYLD_{WT}$, when compared to GFP and $CYLD_{M719V}$, when compared to $CYLD_{WT}$. (**s**) Quantification of the cytoplasmic/nuclear ratio of exogenous TDP-43 shows a marked increase in $CYLD_{WT}$, when compared to cells expressing GFP. The cytoplasmic/nuclear ratio of TDP-43 in $CYLD_{M719V}$-expressing cells was significantly higher than $CYLD_{WT}$. Scale bars = 5 μm. Data is represented as mean ± SEM. a.u. = arbitrary units. *$p<0.05$; **$p<0.01$.
